# Theta activity paradoxically boosts gamma and ripple frequency sensitivity in prefrontal interneurons

**DOI:** 10.1101/2019.12.19.882639

**Authors:** Ricardo Martins Merino, Carolina Leon-Pinzon, Walter Stühmer, Martin Möck, Jochen F. Staiger, Fred Wolf, Andreas Neef

## Abstract

Fast oscillations in cortical circuits critically depend on GABAergic interneurons. Which interneuron types and populations can drive different cortical rhythms, however, remains unresolved and may depend on brain state. Here, we measured the sensitivity of different GABAergic interneurons in prefrontal cortex under conditions mimicking distinct brain states. While fast-spiking neurons always exhibited a wide bandwidth of around 400 Hz, the response properties of spike-frequency adapting interneurons switched with the background input’s statistics. Slowly fluctuating background activity, as typical for sleep or quiet wakefulness, dramatically boosted the neurons’ sensitivity to gamma- and ripple-frequencies. A novel time-resolved dynamic gain analysis revealed rapid sensitivity modulations that enable neurons to periodically boost gamma oscillations and ripples during specific phases of ongoing low-frequency oscillations. This mechanism presumably contributes substantially to cross-frequency coupling and predicts these prefrontal interneurons to be exquisitely sensitive to high-frequency ripples, especially during brain states characterized by slow rhythms.

## Introduction

Collective rhythmic activity is implicated in brain functions from sensory information processing to memory consolidation, often with higher frequency activity bouts locked onto lower frequencies (*1–3*). While the mechanism behind this cross-frequency-coupling is unclear (*3*), the initiation and maintenance of gamma band (30-150 Hz) oscillations are closely associated with fast-spiking (FS), parvalbumin-positive interneurons (*4, 5*). When driven with frequency chirps, and as a result of intrinsic membrane properties, FS neurons fire more robustly at higher input frequencies than spike-frequency adapting (AD), somatostatin-positive interneurons, which are most responsive to lower frequencies (*6*). Nevertheless, recent studies strongly suggest that, under certain conditions, somatostatin-positive interneurons are crucial for gamma oscillations (*7–9*). Could the spectral sensitivity of different interneuron populations perhaps be itself state-dependent? Here we characterized cortical GABAergic interneurons at different *in vivo*-like working points by measuring their dynamic gain (*10–14*).

Dynamic gain quantifies how input in different frequency bands modulates population firing under *in vivo*-like conditions of fluctuating background input. To probe the potential impact of different brain states on spectral sensitivity, we used different types of background inputs that mimic the strength and timescales of correlations in background input across brain states (*15*). We find that both FS and AD interneuron populations can have remarkably wide bandwidths (up to about 500 Hz), making them capable of tracking fast input frequencies well into the range of sharp wave-ripples.

Moreover, our results uncover unanticipated flexibility in AD neurons, which can massively shift their frequency preference, specifically engaging or disengaging with high-frequency rhythms, such as gamma and sharp wave-ripples. The presence or absence of slowly-correlated input drives this sensitivity shift, which can occur within 50 ms, in phase with an ongoing slow rhythm. This observation offers a mechanistic explanation for theta-gamma cross-frequency coupling.

## Results

AD and FS (Figs. 1A and 1B) are the most common firing patterns of somatostatin- and parvalbumin-positive interneurons, respectively (*16*). Their spectral selectivity has been investigated through sub- and supra-threshold cellular responses to simple, purely sinusoidal inputs (*6*) (Figs. 1C and 1D). However, *in vivo*, even when activity on the population level is periodic, the firing of individual neurons appears stochastic, driven by noisy, fluctuating inputs rather than pure sinusoids (*13, 17*). We thus probed the spectral sensitivity of mouse layer 2/3 prefrontal FS and AD interneuron populations under naturalistic operating conditions (Fig. 1E). These dynamic gain measurements require precise control over the neurons’ input to a degree that cannot be attained *in vivo*. We, therefore, used patch-clamp recordings in current-clamp mode in acute prefrontal slices to establish two different regimes of fluctuating input, distinguished by the correlation time τ: the first case, τ = 5 ms, mimics the case of completely asynchronous population activity, when the decay time-constant of synaptic currents is the only source of input correlations (Fig. 1E, black traces). The other input, characterized by a much slower 25 ms correlation time, mimics brain states with population activity exhibiting slow fluctuations, such as quiet wakefulness or slow-wave sleep (*15*) (Fig. 1E, gray traces).

**Fig 1.**
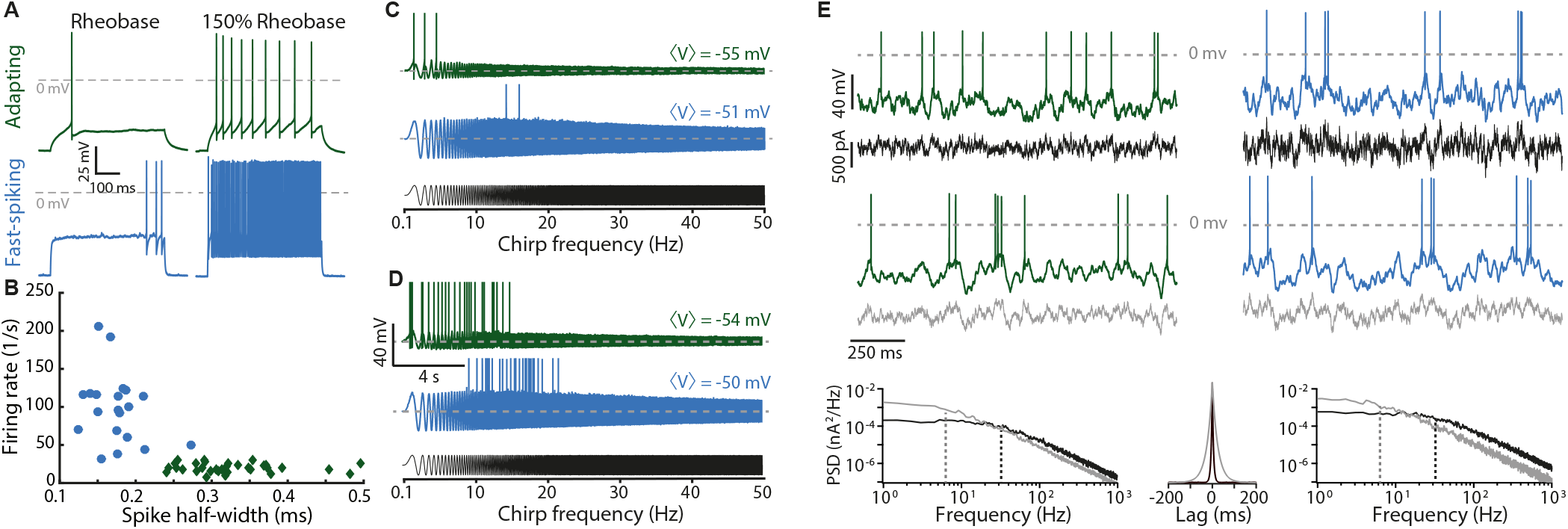
Characterization of neocortical adapting and fast-spiking interneurons. (**A**) Square pulses of 500 ms were used to determine the recorded neuron’s firing pattern at the 150% rheobase level. Shown are representative responses of spike-frequency adapting (AD, green) and fast-spiking (FS, blue) neurons at rheobase and 150 % rheobase. (**B**) Spike half-width and firing rate allow a clear distinction between these cell types. (**C**) Frequency chirp currents (black) have been used to characterize the spectral sensitivity of neurons. They yield action potentials (shown clipped) at lower input frequencies for AD neurons than for FS neurons. (**D**) A slight increase in the offset current, resulting in only a 1 mV depolarization, results in overlapping bandwidths for AD and FS neurons, indicating substantial uncertainty in chirp-based characterizations. (**E**) We assessed neuronal encoding performance in two *in vivo*-like regimes, distinguished by the correlation time of the fluctuating stimuli (τ = 5 ms, black, and τ = 25 ms, gray). The stimulus amplitude at each trial was adjusted to achieve a target operating point (characterized by firing rate and spike train irregularity; see Methods). The corresponding voltage traces of AD and FS neurons are shown above the stimuli, and the power spectral densities (PSDs) and autocorrelations of the inputs are shown at the bottom. The dashed lines in the PSDs indicate the cut-off frequencies (32 Hz and 6.4 Hz) corresponding to the correlation times of the different inputs.

### Input correlations determine frequency selectivity

The spectral sensitivity of interneurons was markedly different from their chirp responses, and for AD cells, it changed drastically between the two conditions (Figure 2A). In the asynchronous regime, AD neurons respond preferentially to slow components, with the highest sensitivity in the 2-4 Hz range (mean dynamic gain = 119 Hz/nA, 95% bootstrap confidence interval: [117, 122]). The average gain in the gamma range (Fig. 2A, shaded region) reaches only 62% of the average at lower frequencies (< 20 Hz) (64 Hz/nA [63, 65] vs 103 Hz/nA [101, 105]). These values mean that the addition of a small, 10 pA sinusoidal modulation (equivalent in magnitude to a single synaptic event) on top of the irregularly fluctuating background input would modulate the AD population’s firing rate by 1.2 Hz in response to a superimposed 3 Hz input, but it would modulate the firing rate only by 0.6 Hz for 60 Hz, indicating a clear preference for lower frequencies. This preference, however, changed completely when AD neurons were exposed to slowly fluctuating input such that their preferred frequency shifted from 3 Hz to 200 Hz. The gain at 2-4 Hz dropped from 119 Hz/nA [117, 122] to 91 Hz/nA [89, 92], and the gain at 200 Hz increased from 74 Hz/nA [72, 76] to 119 Hz/nA [113, 124]. Figure 2B demonstrates the occurrence of this shift in two individual AD neurons. With this abrupt change in frequency preference, AD neurons in the synchronous regime become more sensitive to gamma input than to lower frequencies (average gains: 97 Hz/nA [94, 101] vs 81 Hz/nA [80, 83]). Altogether, these data reveal that, during network states characterized by slow background fluctuations, AD cells tune themselves to gamma and higher-frequency input. FS interneurons, on the other hand, preferentially transmit high frequencies irrespective of the input correlations, with a maximum sensitivity around 200-250 Hz (Fig. 2C). Both, FS interneurons and, given sufficiently slow input components, AD interneurons have a remarkably wide bandwidth, with a high-frequency limit well above 400 Hz, an order of magnitude higher than expected from their chirp-responses.

**Fig. 2.**
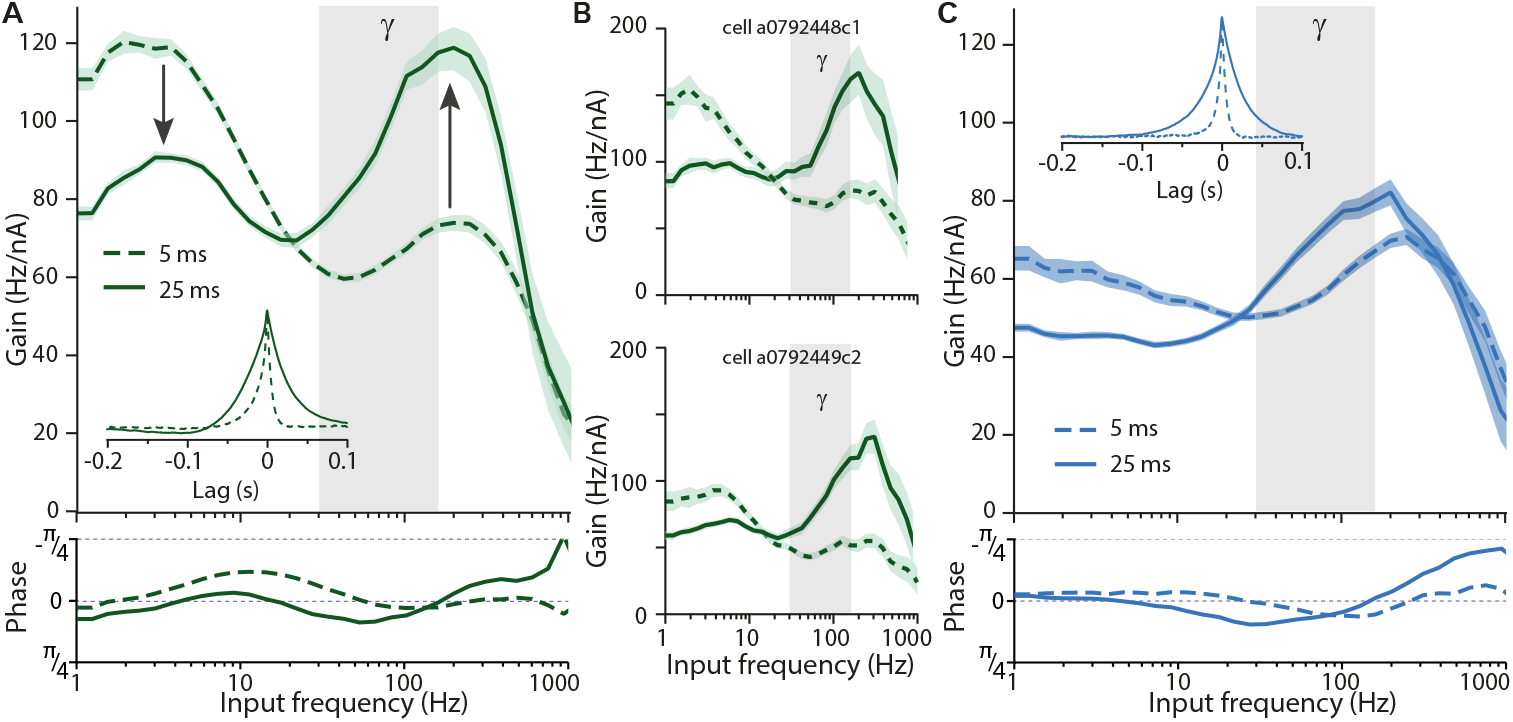
Spectral selectivity of AD neurons drastically shifts for different background fluctuations. (**A**) (top) Gain of AD neurons tested with inputs with two different correlation times. Under fast background input, τ = 5 ms (dashed lines, n = 12), AD neurons modulate their firing rate strongest in response to lower frequencies. Under slow background, τ = 25 ms (continuous line, n = 10), the frequency preference shifts (arrows) and the firing rate is modulated mainly by high frequencies. Mean firing rate and coefficient of variation of the interspike intervals were 4.0 Hz ± 0.2 and 0.99 ± 0.02 (5 ms input) and 3.7 Hz ± 0.2 and 1.03 ± 0.05 (25 ms input), respectively. Inset: spike-triggered average input across all recorded cells tested with the same correlation times. Gains were calculated by taking the ratio of the Fourier transforms of the spike-triggered average and of the autocorrelation function of the input. Gray columns represent the gamma frequency band, and the shaded region around gain curves represents the 95% bootstrap confidence interval. (Bottom) Phase of firing rate modulation with respect to input. No substantial phase-delays are associated with action potential generation. (**B**) Individual gain curves of two AD cells from **A** for both correlation times. The drastic shift in frequency preference is clearly visible at the single-cell level. (**C**) As in **A**, but for FS neurons. Those display a wide bandwidth and no drastic shifts in frequency preference (τ = 5 ms, dashed lines, n = 7; τ = 25 ms, continuous lines, n = 9). Grand-averaged firing rate and coefficient of variation of the interspike intervals were 5.0 Hz ± 0.6 and 1.12 ± 0.24 (5 ms) and 3.6 Hz ± 0.2 and 1.47 ± 0.10 (25 ms). Numbers are given as mean ± SEM.

### Theta input reliably boosts gamma- and ripple-sensitivity of spike-frequency adapting neurons

This input-dependent spectral sensitivity might allow AD neurons to provide brain-state-dependent feedback input into the local cortical circuit. Interestingly, AD neurons shift their preference to the gamma-band when lower frequencies dominate their input. This suggests that the presence of theta oscillations (4-12 Hz) could tune them to higher frequencies, boosting gamma components. To test this hypothesis, we exposed AD neurons to the rapidly fluctuating, 5 ms correlated background input, either on its own or supplemented with theta-band components (Fig. 3A). We found that the theta-band components indeed boosted the gain for frequencies above 30 Hz, with the average gamma-band gain increasing from 71 Hz/nA [69, 73] to 86 Hz/nA [83, 89] and the average gain in the ripple-band increasing from 79 Hz/nA [77, 82] to 105 Hz/nA [101, 110] (Figure 3B). The gamma/theta ratio increased in 9 out of 10 cells, from 0.56 ± 0.02 to 0.74 ± 0.05, while the ripple/theta ratio increased from 0.63 ± 0.02 to 0.88 ± 0.05 (mean ± SEM; Fig. 3C), revealing that, indeed, an increase in theta power boosts gamma- and ripple-sensitivity of AD neurons.

**Figure 3.**
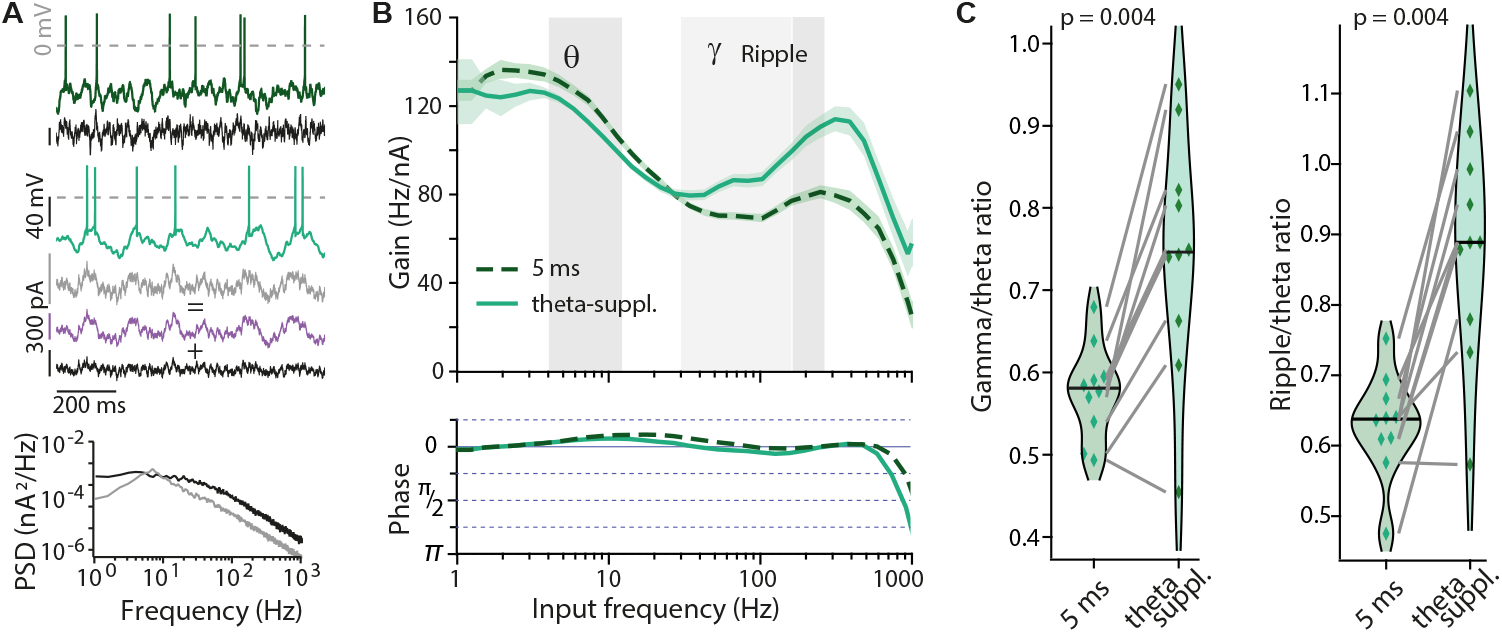
Increasing theta input to AD neurons boosts sensitivity to gamma and ripple frequencies. (**A**) Sample stimuli and voltage traces (dark green, 5 ms input; light green, theta-supplemented input) and power spectral density of noisy inputs with τ = 5 ms (black) and theta-supplemented 5 ms input (gray). Theta-supplemented input was constructed by adding a theta bandpass filtered white noise input (purple) to the 5 ms input. Grand-averaged firing rate and coefficient of variation of the interspike intervals were 4.58 Hz ± 0.23 and 0.97 ± 0.03 (5 ms input) and 4.78 Hz ± 0.26 and 0.92 ± 0.04 (theta-supplemented 5 ms input). (**B**) (top) Gain of AD cells tested with both, τ = 5 ms (dashed line) and theta-supplemented 5 ms (continuous line) inputs (n = 10). Boosting theta in the input paradoxically reduces the sensitivity of AD neurons to this frequency band while promoting sensitivity to gamma and ripple frequencies (150-250Hz). (Bottom) The phase of firing rate modulation with respect to the input. No substantial phase-delays are associated with action potential generation, even though the gain magnitude is modulated. (**C**) Ratios between average gains at gamma and theta (left) and ripple and theta (right) for the individual neurons (diamonds). Both ratios increase when theta power in the input is increased. Gamma/theta ratio for τ = 5 ms: 0.56 ± 0.02 vs theta-supplemented 5 ms: 0.74 ± 0.05, n = 10 (paired sample, two-sided Wilcoxon signed-rank test, W = 1, p = 0.004). Ripple/theta ratio for τ = 5 ms: 0.63 ± 0.02 vs theta-supplemented 5 ms: 0.88 ± 0.05, n = 10 (two-sided Wilcoxon signed-rank test, W = 0, p = 0.002). Violin plots show the medians as bars. Numbers are given as mean ± SEM

### Theta phase determines gamma sensitivity

The dynamic gain curves above are based on the timing of all action potentials (APs) fired during a long stimulus period. They represent the average frequency selectivity for a given input statistics and allowed us to detect the boosting for gamma- and ripple frequencies during 30 second long periods of theta-dominated input. During *in vivo* activity, however, short gamma-bursts or ripples occur phase-locked to slower rhythms, consistent with the idea that neurons might be recruited to high-frequency rhythms within a few dozen milliseconds. Specifically, theta-gamma cross-frequency coupling suggests a modulation of gamma sensitivity throughout the phase of the ongoing theta rhythm. To test whether AD neurons indeed display such a modulation, we developed a time-resolved decomposition of the dynamic gain. To this end, we reanalyzing the data obtained with the theta-supplemented stimulus (Fig. 3), we determined the phase *φ_θ_* of the stimulus’ theta band at each AP time. We sorted the APs into three groups, according to *φ_θ_*. The boundaries between groups, *φ_θ_* = −0.017 and *φ_θ_* = 0.476, were chosen so that each group contained one-third of all APs (Fig. 4A and B, Methods). For each group, the average gain across all ten neurons was determined as before (Fig. 4C). As expected for a meaningful decomposition, the three gain components combined to replicate the overall dynamic gain (Inset in fig. 4C). For each neuron, we calculated three dynamic gain curves, each derived from all the APs belonging to one *φ_θ_* group. The neuron’s ability to lock to gamma rhythms was quantified as the average gain value in the gamma range (30-150Hz). Across the ten neurons, the median value of this gamma sensitivity increased from 27.4 Hz/nA to 37.5 Hz/nA as *φ_θ_* goes from the first to the third group, revealing a theta-phase dependent locking of APs to gamma inputs (Fig. 4D). The substantially increased gamma sensitivity for APs fired later during a theta cycle indicates that AD neurons’ frequency tuning changes within a quarter of a theta cycle, i.e., within less than 50 ms.

**Fig. 4.**
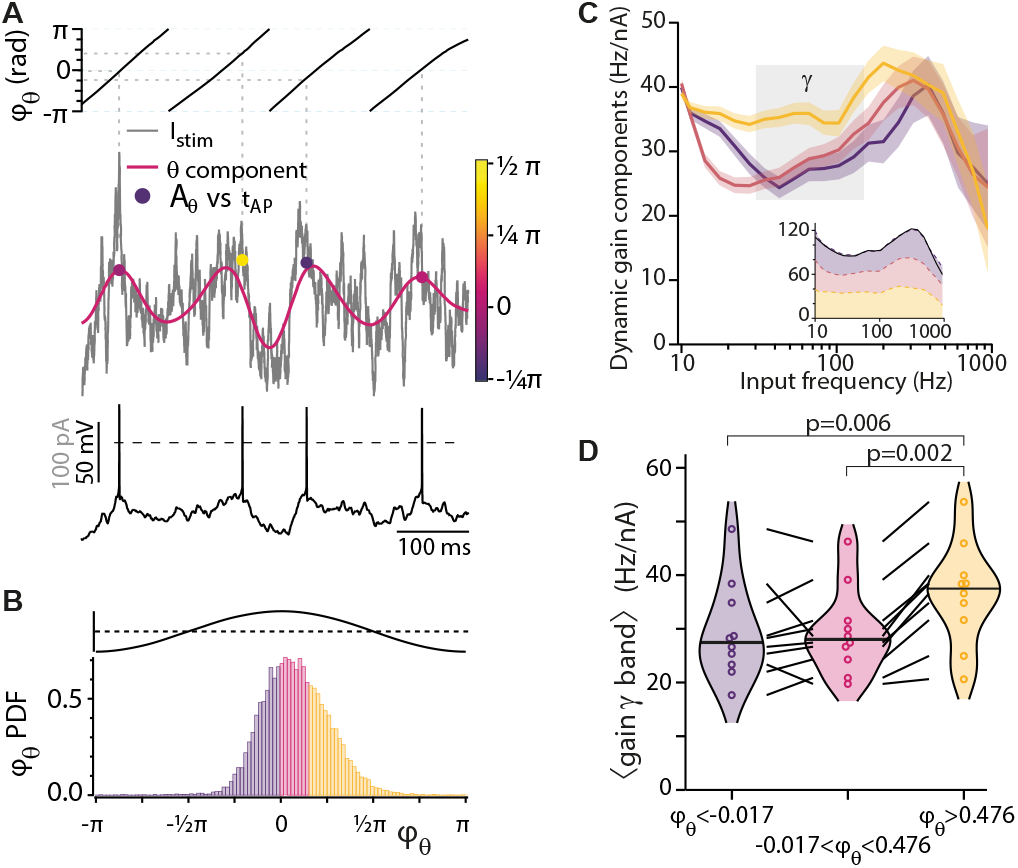
Sensitivity to gamma frequencies is modulated during progression through the theta cycle. (**A**) Analysis of theta components showing the membrane voltage (bottom, black), input current (grey), its theta component (magenta, see Methods), and points indicating the time of action potentials, plotted against the instantaneous theta amplitude A_θ_ obtained by Hilbert-Analysis. The color code corresponds to the theta phase (φ_θ_, top). (**B**) A probability density plot of φ_θ_ with three differently colored phase intervals. Each interval contains one-third of all APs. The top trace indicates the cosine relation between the theta component’s phase and amplitude. (**C**) The dynamic gain curves of the three components (color code as in B) have distinctive frequency dependencies. Their 95% confidence intervals (shaded) separate in the beta (12 – 30 Hz) and gamma frequency bands. In the inset, the three components are stacked. Their sum closely reproduces the overall dynamic gain obtained with the theta-supplemented input from fig. 3 B (re-ploted in black in the inset) (**D**) AD neurons’ sensitivity to gamma frequencies (Methods) increases during the theta cycle from the lower two φ_θ_-intervals: 29.4 ± 2.9 Hz/nA (purple, left) and 29.4 ± 2.6 Hz/nA (magenta, center), to the APs with the highest φ_θ_ values: 36.5 ± 3.0 Hz/nA (orange, right). Gamma sensitivity in the latest interval (orange) is significantly higher than in the first (purple, W=2, p=0.006) and the second (magenta, W=0, p=0.002), based on paired sample, two-sided Wilcoxon signed-rank tests.

## Discussion

Our study reveals a novel type of dynamic regulation of AD neuron’s response properties. When the neurons’ input fluctuates rapidly, as during active wakefulness, our data support the traditional picture in which AD neurons preferentially encode low-frequency input, and FS neurons encode high-frequency (>30 Hz) input. A drastic change in the frequency preference of AD neurons occurs, however, when input correlations are slow, as during brain states featuring low-frequency dominated local field potentials. Under such conditions, AD neurons react preferentially to gamma and ripple frequencies. Our findings thus uncover an unanticipated flexibility of interneuron function that allows brain states to tune AD neuron population coding and might underlie their reported contribution to gamma oscillations (*7, 9*).

Previously, dynamic gain curves were studied as essentially static properties, determined from minute-long stimuli of stationary stochastic properties. Our time-resolved analysis revealed that AD neurons rapidly respond to fluctuations in input statistics, increasing their dynamic gain in the gamma-band by 50% within a few dozen milliseconds. Our decomposition approach provides a powerful extension to current population encoding analyses. It allows, for instance, a quantitative comparison between the encoding capacity of APs within ripples vs outside ripples, or of AP duplets as compared to isolated APs.

Increased encoding of high frequencies (>30 Hz) during particular phases of strong, slow (theta) components (Figs. 3, 4) closely matches the phenomenon of theta-gamma cross-frequency coupling (*18*). In the dynamics of recurrent local circuits, the dynamic gain is a main component to the feedback gain that determines whether a collective oscillation is amplified or dies out. In theoretical studies of population oscillations in synchronous (*2*) or asynchronous (*19*) network states, the magnitude and the phase of the dynamic gain are key determinants of oscillations strength and frequency (*20*). Therefore, the small phase-delays associated with AP generation (Figs. 2A, 2C and 3B) and the input-dependent increase in gain magnitude predict a boost of gamma oscillation amplitude in the presence of theta-frequency input components and in particular during late theta-phases. The dynamic tuning of spectral sensitivity in phase with slow input fluctuations is, to our knowledge, the first mechanism coupling gamma amplitude to theta oscillations that is based on cellular electrophysiological properties.

The wide bandwidth of AD and FS neurons of up to 500 Hz and a maximal sensitivity reached around 200 Hz is by itself a striking phenomenon. In cortical pyramidal neurons, high bandwidth dynamic gain is known to mediate the sub-millisecond precision of population coding for input changes (*21*), but what function could a narrow preference band at around 200 Hz serve? Retrieval and consolidation of episodic memory require a complex and precise replay of activity by cell assemblies in the form of high-frequency sharp wave-ripples (150-250 Hz). Intriguingly, these occur specifically during periods of synchronous network activity, such as during slow-wave sleep or quiet wakefulness (*22*), when, as we showed, AD and FS neurons are most sensitive to high frequencies. Given the input-dependent selectivity switch in AD neurons, slow oscillations may, in general, boost high-frequency sensitivity of interneurons and specifically allow AD neurons to tune in to ripple-related inputs and disinhibit cortical circuits in a precisely timed manner.

## Materials and Methods

### Animals and slice preparation

All experiments were performed in accordance with institutional and state regulations (Niedersächsisches Landesamt für Verbraucherschutz und Lebensmittelsicherheit). Experiments were performed in 3 to 8-week-old mice of either sex from five different mouse lines. Two lines target mostly AD interneurons: GIN (FVB-Tg(GadGFP)45704Swn, The Jackson Lab #003718) and SOMCrexAi9 (Ssttm2.1(cre)Zjh/J, The Jackson Lab #013044, crossbred with B6.Cg-GT(ROSA)26Sor^tm9(CAG-tdTomato)Hze/J, The Jackson Lab #007909); and three lines target mostly FS interneurons: PVCre (*23*), PVCrexAi32 (PVCre crossbred with B6;129S-Gt(ROSA)26Sortm32(CAG-COP4*H134R/EYFP)Hze/J, The Jackson Lab # 012569), and Nkx2.1CreERxAi14 (Nkx2-1tm1.1(cre/ERT2)Zjh/J, The Jackson Lab # 014552, crossbred with B6;129S6-Gt(ROSA)26Sortm14(CAG-tdTomato)Hze/J, The Jackson Lab # 007908). Animals were kept in standard 12h light regime with water and food *ad libidum*. Animals were intraperitoneally-injected with a mixture of ketamine and xylazine in PBS (respectively 100 and 20 mg/kg of body weight) and decapitated. The brain was quickly removed and kept in ice-cold, carbogen-saturated cutting solution containing, in mM, 125 NaCl, 2.5 KCl, 26 NaHCO_3_, 1.25 NaH_2_PO_4_, 0.4 Ascorbic Acid, 4 Na-Lactate, 25 Glucose, 1 MgCl_2_, 2 CaCl_2_ (~315 mOsm, pH 7.4). 300-μm-thick coronal neocortical slices were made in a VT1200S Vibratome (Leica) and incubated at 35°C in carbogen-saturated recording solution (aCSF, in mM: 125 NaCl, 4 KCl, 26 NaHCO_3_, 10 glucose, 1.3 MgCl_2_, 2 CaCl_2_) until recorded.

### Patch-clamp recordings

One slice at a time was transferred to a heated recording chamber (PH6 and RC-27L, Warner Instruments) and mechanically stabilized with a slice hold-down (SHD-27LH/15, Warner Instruments). Throughout the experiment, the slice was gravitationally perfused with warm aCSF through an in-line heater (HPT-2, Alasciences) at a 1-2.5 ml/min flow rate. Both the recording chamber and the in-line heater were controlled by a TC-20 temperature controller (NPI electronics). Temperature settings were adjusted so that a thermistor measured a target temperature of 36 ± 1° C at the slice position. Slices were visualized in an Axio Examiner.D1 microscope (Zeiss) equipped with a W Plan-Apochromat 40x/1.0 DIC objective. Cells were visualized with infrared differential interference contrast optics (Zeiss), and fluorescent signal was imaged with a multi-wavelength LED source (pE-4000, CoolLed) and a CCD camera (MD061RU-SY, Ximea). 4-6 MOhm pipettes were prepared from borosilicate glass capillaries (PG10165-4, World Precision Instruments) in a vertical puller (PIP 6, HEKA). Internal solution contained, in mM, 135 K-Gluconate, 10 KCl, 4 NaCl, 0.1 Na_4_EGTA, 1 Mg-ATP, 0.3 Na-GTP, 10 Hepes, 0.5 Na_2_-Phosphocreatine and 0.2% (w/v) biocytin (285-290 MOsm, pH adjusted to 7.3). Whole-cell current-clamp recordings were made in an EPC-10 Double amplifier controlled by Patchmaster (Heka). Fast and slow capacitances and series resistance were carefully adjusted in voltage-clamp mode before recording; fast capacitances while in on-cell configuration, slow capacitances after achieving whole-cell configuration. Series resistance was 90-100% compensated with a feedback time constant of 10 μs. Voltage signals were low-pass filtered at 8.8 kHz and digitized at 100 kHz. Data analyses were performed in custom-written Matlab 2014b (Mathworks) and Igor Pro 8 (Wavemetrics) programs. Liquid junction potential of −14 mV was not corrected. All experiments were performed in the presence of blockers of GABA receptors (picrotoxin, 30 μM, Sigma) and glutamate receptors (NBQX, 10 μM, Tocris; and DL-AP5, 30 μM, Sigma).

### Characterization of action potential firing patterns

Layer 2/3 interneurons were identified via fluorescence imaging. In order to identify their electrical type, 500-ms-long current steps were applied. Current amplitude was increased in 15 pA steps until at least 1.5 times rheobase, the level at which the characterization of the firing pattern was made. Only cells exhibiting clear fast-spiking (including stuttering cells) and adapting electrical types were included in the analysis.

### Dynamic gain calculation

Population frequency-response characterization was restricted to layer 2/3 prefrontal FS and AD interneurons and was assessed as previously described (*11, 12*). This analysis aims to achieve an *in vivo*-like operating point, mimicking a situation in which a high rate of synaptic inputs provides a continuously changing net background current, and a neurons’ firing is driven not by the average input but by its transient depolarizing excursions (*17*). Fluctuating current inputs were synthesized as Ornstein-Uhlenbeck noises *x(t)* with either 5 or 25 ms correlation time. These values were chosen to approximate the case of uncorrelated inputs filtered through the synaptic currents’ decay time-constants (5 ms) or the case of slow temporal correlations in the input due to correlated network activity (25 ms). Inputs’ standard deviation was adjusted to obtain similar firing rates (around 4 Hz) and coefficients of variation of the interspike intervals (around 1) for both correlation times. Neurons were first depolarized to −60 mV with DC current and different realizations of the fluctuating noise were injected in 30-s-long episodes, separated by 15-s-long resting, for as long as the recording did not display signs of deterioration, such as baseline drifts or spike overshooting to positive voltages less than 20 mV. For experiments presented in figure 3, a theta-power enhanced stimulus was created by adding a 4-12 Hz bandpass filtered white noise to the 5 ms input. The standard deviation of the bandpass filtered signal was normalized to two times the standard deviation of the 5 ms signal. APs were detected as 0 mV crossings on the voltage trace and the AP times were annotated. From these, a spike-triggered average input current (STA) was obtained by summing up 1-s-long stimulus segments centered on the AP times for all cells of a given condition and dividing by the total number of APs.

The complex dynamic gain function *G(f)* was calculated as the ratio of the Fourier transform of the STA, *F*|STA|, and the Fourier transform of the autocorrelation of the stimulus, *F*|c_ss_(τ)| where

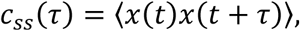

and *τ* denotes the time lag. In order to improve signal-to-noise ratio, *G(f)* was filtered in the frequency domain by a Gaussian filter *w(f’)* centered at frequency *f’=f* and a frequency-dependent window size with standard deviation of *f/2π*.

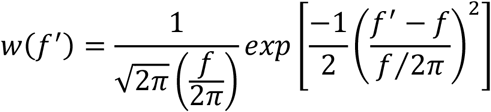

The filtered dynamic gain function *G_w_(f)* thus becomes

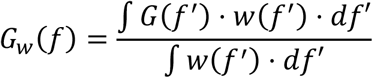

The magnitude and phase of this filtered, complex dynamic gain function are reported in figures 2 and 3. For the dynamic gains reported in figure 2, the data comprise of: for AD neurons, 19563 spikes from 12 cells and 20427 spikes from 10 cells (5 ms and 25 ms respectively), and, for FS neurons, 9792 spikes from 7 cells and 15023 spikes from 9 cells (5 ms and 25 ms, respectively). Five of the 10 AD neurons tested with 5 ms-correlated stimuli were also tested with the theta-supplemented 5 ms input. In addition to these, another 5 were used to obtain the gains in figure 3 (14847 spikes, for 5 ms stimulus and 18067 spikes for theta-supplemented 5 ms stimulus). Confidence intervals were obtained by bootstrap resampling. 2000 bootstrapped gain curves were calculated from the same number of STAs obtained by randomly sampling from all STAs used in the population gain calculation. The confidence intervals are defined as the 2.5^th^ and 97.5^th^ percentiles at each frequency point in the 2000 gain curves. The distribution of this bootstrap statistic was not different from normal (Kolmogorov-Smirnov test). To identify the portions of the gain curves that are significantly different from zero, we calculated a noise floor. It was calculated by cyclically shifting original spike times by a random time interval, larger than 5 correlation times, and calculating 2000 “random time-triggered averages”, which were used to calculate “gain curves”. The noise floor was defined as the 95^th^ percentile of these “gain curves”. The gain curves in figures 2 and 3 were displayed either until they were crossed by the noise floor or up to 1000 Hz, if noise floor crossing happened at a frequency > 1000 Hz.

### Hilbert Analysis

The stimulus’ theta phase component was extracted by filtering with an infinite impulse response bandpass filter with 6 pole Butterworth characteristics. The filter was designed in Igor Pro 8 with pass-band limits of 3.5 and 10.5 Hz for the sample time of 100kHz. Used twice, once in forward time, once in reversed time, the filter results in zero-delay filtering of the input. Fourier analysis of the input and output shows effective isolation of the 4 to 12 Hz component. The phase and amplitude of this component were obtained by conventional Hilbert analysis. APs were stratified according to the phase of the theta component at the AP time to perform the analysis in figure 4.

### Statistics

Paired samples, two-sided Wilcoxon rank tests were performed to test the single neuron data in figures 3 and 4. W-statistics and p values are given in figures and legends. The p-values in figure 4 are not corrected for the dual comparison.

## Funding

Bundesministerium für Bildung und Forschung (BMBF, Federal Ministry of Education and Research) grant 01GQ1005B (FW, AN)

Bundesministerium für Bildung und Forschung (BMBF, Federal Ministry of Education and Research) grant 01GQ1005E (FW)

Deutsche Forschungsgemeinschaft (DFG, German Research Foundation) – 436260547, in relation to NeuroNex (NSF 2015276)

VW Foundation grant ZN2632 (FW)

GGNB Excellence Stipend of the University of Göttingen (RMM)

Max Planck Society

## Author contributions

R.M.M., C.L.P., F.W., and A.N. conceived the study. R.M.M. performed the experiments with contributions from M.M. R.M.M., C.L.P., F.W., and A.N. analyzed and interpreted the data. F.W., A.N., W.S., and J.F.S. provided resources. All authors discussed and interpreted the data. R.M.M., F.W, and A.N. wrote the paper with inputs from all other authors.

## Data and materials availability

All data are available from the corresponding author upon reasonable request. Raw data underlying the dynamic gain curves can be downloaded from this permanent repository at the Max Planck Digital Library: https://edmond.mpdl.mpg.de/imeji/collection/pdxNFpqJurbDDeop. *This permalink is for review purposes. It will be replaced with a DOI*. The code, written in IgorPro 8.0 and 9.0, used to analyze raw data and generate the dynamic gain curves, is included in the data repository. The code is continuously maintained. The latest version is available at https://github.com/Anneef/AnTools.

